# Genetic correlations across genetically determined and developmentally plastic alternative reproductive tactics

**DOI:** 10.1101/768721

**Authors:** Jessica K. Abbott, Oscar Rios-Cardenas, Molly Morris

## Abstract

Alternative reproductive tactics occur when individuals of the same sex have a suite of morphological and/or behavioural traits that allow them to pursue different reproductive strategies. A common pattern is e.g. the existence of “courter” and “sneaker” tactics within males. We have previously argued that alternative reproductive tactics should be subject to genetic conflict over the phenotypic expression of traits, similar to sexual antagonism. In this process, which we called intra-locus tactical conflict, genetically determined tactics experience conflicting selection on a shared phenotypic trait, such as body size, but a positive genetic correlation between tactics in body size prevents either tactic from reaching its optimum. Recently, other authors have attempted to extend this idea to developmentally plastic alternative reproductive tactics, with mixed results. However, it is not clear whether we should expect intra-locus tactical conflict in developmentally plastic tactics or not. We have therefore run a series of simulation models investigating under what conditions we should expect to see positive estimates of the inter-tactical genetic correlation, since a positive genetic correlation is a prerequisite for the existence of intra-locus tactical conflict. We found that for autosomal, X-linked, and Y-linked genetically-determined tactics, estimated inter-tactical genetic correlations were generally high. However, for developmentally plastic tactics, the genetic correlation depends on the properties of the switching threshold between tactics. If it is fixed, then estimated genetic correlations are positive, but if there is genetic variation in the switch-point, then any sign and magnitude of estimated genetic correlation is possible, even for highly heritable traits where the true underlying correlation is perfect. This means that caution should be used when investigating genetic constraints in plastic phenotypes.

## Introduction

Alternative reproductive tactics (ARTs) are different ways of achieving reproductive success within a sex, and often involve suites of behavioral, morphological and life history traits. ARTs are expected to evolve when sexual selection is strong and multiple strategies are possible (reviewed in Shuster and Wade 2003). Studies of ARTs across a range of taxa (reviewed in Oliveira et al. 2008) have led to a better understanding of the maintenance of genetic variation given strong sexual selection, as well the adaptive nature of that variation (i.e. the best phenotype for using one tactic is often not the best phenotype for a different tactic). The discontinuous expression of at least one or more traits in either males or females is often the first indication of an ART (e.g. Gross 1996; Brockmann 2001). However, there is a growing appreciation for the role of intralocus tactical conflict (IATC) in constraining the evolution of differences between the ARTs (tactical dimorphism), such that even traits that are more or less continuous within a sex may have more than one underlying optimum (Abbott et al. 2019).

Individuals from different ARTs will share many homologous traits, however if the optimal state for these traits differ depending on the ART, this will lead to opposing selection (i.e. tactically disruptive selection). When traits that are genetically correlated across the ARTs are not at their adaptive optimum, tactically disruptive selection can generate intralocus tactical conflict (Morris et al. 2013; Buzatto et al. 2015). Studies of intralocus tactical conflict have the potential to increase our awareness of cases where expressed states are not necessarily optimal due to the evolutionary constraints (e.g. slower growth rates, a lack of behavioural plasticity; Abbott et al. 2019). IATC also has the potential to lead to a better understanding of the role of ecological variation across populations in producing the patterns of divergence between ARTs, as well as provide us with a better understanding of the relationship between ARTs and rapid speciation (Abbott et al. 2019). The criteria for demonstrating intralocus tactical conflict include a positive genetic correlation between the ARTs, detecting different optima for the trait across ARTs, and evidence that the ARTs are not at their optima for the trait (Morris et al. 2013).

Inter-tactical genetic correlations measure the extent of similarity between the additive effects of alleles when expressed in different tactics. The ideal method for determining inter-tactical genetic correlations when ARTs are genetically fixed is a multigeneration half-sib breeding design, followed by the statistical decomposition of the genetic variance into its many different components (Falconer and Mackay 1996). Given the prevalence of genetic correlations across the sexes even when sexual dimorphism has evolved (e.g. Harano et al. 2010; Poissant et al. 2010), it can be assumed that these correlations are not temporary or transitional stages, highlighting the importance of their estimation across ARTs as well as across the sexes. By current estimations, most ARTs are developmentally plastic (Oliveira et al. 2008); however, this consensus may change as the studies of the proximal mechanisms behind these ARTs increase.

West-Eberhard (1986) described how the loss of an alternative phenotype could play a role in speciation through the release “from constraints of having to accommodate multiple alternatives” (pg 1388) within a shared genome. And yet, the idea that developmental plasticity decouples the development of the alternative morphs, allowing them to evolve independently, is prevalent in the literature (reviewed in Tomkins and Hazel 2007). Therefore, a better understanding of the potential for genetic correlations between ARTs that are both genetically fixed and developmentally plastic is needed to determine the extent to which intralocus tactical conflict may be influencing the evolution of ARTs. There are a few empirical studies that have examined genetic correlations across developmentally plastic ARTs. Considering male traits across two species with tactically dimorphic male ARTs that are environmentally influenced, Pike et al. (2017), detected very weak genetic correlations in one species (earwigs, *Forficula auricularia*), and significant correlations in another species (acarid mites, *Rhizoglyphus echinopus*). The genetic correlations in the acarid mite have been further confirmed through artificial selection experiments (Buzatto et al. 2018).

Genetically-determined ARTs can be divided into two types that could potentially differ in their propensity for genetic correlations. First, allelic variation at autosomal loci can influence polymorphisms, as in a marine isopod (Shuster and Wade 1991). Second, genetic polymorphisms may be correlated with the genes influencing sex, as in the swordtail fishes where male ARTs have been linked to genetic variation on the Y-chromosome (Zimmerer and Kallman 1989; Lampert et al. 2010). Developmentally plastic ARTs, on the other hand, have been proposed to evolve via environmental thresholds. In this model, the environment influences alternative phenotypes through a genetically-determined threshold (Hazel et al. 1990; Hutchings and Myers 1994), which may be able to respond rapidly to artificial selection (Emlen 1996). Studies of several species have provided evidence for the environmental threshold model of developmentally plastic ARTs: Atlantic salmon, where males can either mature sexually early in life in freshwater or more commonly only after completing a migration at sea (Lepais et al. 2017); horned beetles, where some males develop horns and some do not (Emlen 1996); and bulb mites (Buzatto et al. 2015). In the current study, we use simulations to examine the potential for genetic correlations across ARTs that vary in their underlying mechanisms, including an autosomal ART locus, an X- or Y-linked ART locus, a fixed threshold for plastically determined tactics, and genetic variation in the switching threshold between plastic tactics.

Intertactical additive genetic correlations, similar to *r*_mf_ between the sexes (Lande 1980), are predictive of the potential for future independent evolution of the ARTs within a population. Here we considered the genetic correlation of traits across ARTs that are not directly linked to the locus (or loci) producing the ART. For example, in the case of the autosomal supergene producing differences across male ARTs in ruffs (Küpper et al. 2015), the inversion has linked a suite of traits involved in the ARTs. However, by considering the extent to which these correlations will be present for traits not directly linked to the ARTs, we can determine the extent to which IATC can influence and potentially constrain their evolution. Our results suggest that for autosomal, X-linked, and Y-linked genetically-determined tactics, estimated genetic correlations are, as expected, generally high. However, for developmentally plastic tactics, the estimated genetic correlation depends on the properties of the switching threshold between tactics, such that genetic variation in the switching threshold can lead to a range of estimates even when the true underlying genetic correlation is perfect.

## Simulation model

In the case of genetically-determined tactics, all simulations assume a single ART locus with two alternative alleles producing different alternative reproductive tactics. If tactic is developmentally plastic, then the phenotype determines the threshold for switching between tactics. Different relationships between the phenotype and the threshold are considered (see below). The trait that can experience tactical antagonism is a quantitative trait determined by many loci that are spread throughout the genome (e.g. body size). This means that there is no effect of linkage between the ART locus and the trait.

The simulations are designed in such a way that they resemble the simplest form of an animal model:

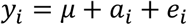

Where *y_i_* is the phenotypic trait value of individual *i, μ* is the phenotypic mean in the population, *a_i_* is the breeding value, and *e_i_* is the residual error. Parameters in the simulations are therefore the number of families (fixed at 1000), the population mean (arbitrarily fixed at 50), the population standard deviation (used in randomly generating a breeding value for each family; arbitrarily fixed at 5 unless otherwise stated), the within-family standard deviation (used in generating developmental noise for each individual, and equal to the residual standard error), the magnitude of the trait difference between the ARTs (arbitrarily fixed at 5 unless otherwise stated), and the offspring number (arbitrarily fixed at 10 unless otherwise stated). Heritability measured as *V_a_/V_p_* is therefore equal to the population SD/(population SD + within-family SD). For developmentally plastic tactics there is an additional parameter for the switching threshold. The inter-tactical correlation was estimated from within-family means, as in Buzatto et al. (2015). For Y-linked tactics, we also included an analysis using MCMCglmm (see below). Datasets were simulated 1000 times each, in order to obtain information about stochastic variation in correlation estimates for a given set of parameter values.

We examine several different scenarios, including an autosomal ART locus, an X- or Y-linked ART locus, a fixed threshold for switching tactics, and various ways of relating the switching threshold to the breeding value. Note that in all cases the underlying genetic architecture of the quantitative trait is a perfect genetic correlation between tactics, so what we investigate with our simulations is variation in the empirical estimate of the genetic correlation, and how well it corresponds to the true genetic architecture.

## Autosomal ART locus

This is the simplest possible case, and was simply analysed to provide a baseline for comparison with the other scenarios. We assume that at least some families are capable of producing a mix of different ARTs, for example when two heterozygotes for a dominant ART locus mate with each other and produce offspring of all possible genotypes. We varied the population standard deviation, the within-family standard deviation, offspring number, and the mean trait difference between ARTs, and investigated heritability and the magnitude of the inter-tactical genetic correlation.

We found that, as expected, estimated inter-tactical correlations were high for an autosomal ART locus, and increased with increasing heritability (Figure 1). This is a logical result; if the variation within a family is as large as (or larger than) the variation in breeding values across the entire population, it will be more difficult to detect the inter-tactical correlation. Interestingly, we could also show that the estimate of the inter-tactical genetic correlation was often higher than the heritability of the trait (Figure 1, dashed line shows 1:1 relationship), which suggests that the underlying genetic architecture is more important in determining the estimated genetic correlation than the observed heritability. Increasing or decreasing the trait difference between the tactics does not affect these conclusions, but assigning different levels of developmental noise to the two ARTs, or reducing family size, both decrease the estimate of the correlation (Figure 1). This is also an intuitive result since there will be more error in the estimate when the family size is low. The effects of trait difference between tactics and family size is consistent across all other scenarios discussed below (data not shown).

**Figure 1:**
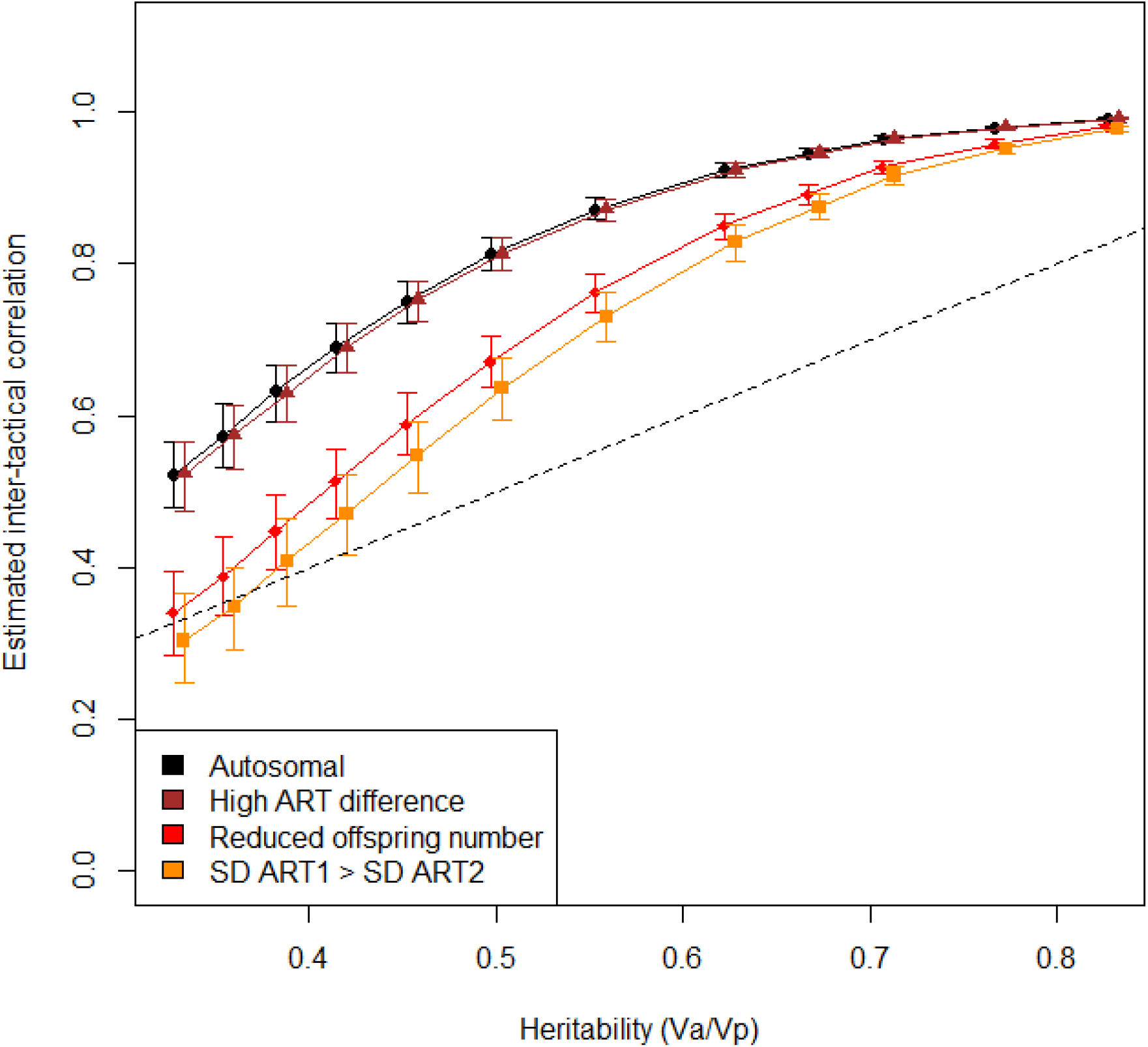
Estimated inter-tactical genetic correlation for a quantitative trait dependent on its heritability, in the case of an autosomal ART locus. Error bars denote 95% confidence intervals and points are jittered to avoid overlap. Estimates increase with increasing heritability. Increasing the magnitude of the difference between ARTs (here by a factor of two) does not affect the estimated genetic correlation. However, reducing the offspring number within each family (by 50% shown here) and a higher variance in one ART (SD ART1 = 2*SD ART2 shown here) both decrease the magnitude of the estimated genetic correlation. Points falling above the dashed 1:1 line have a higher estimated genetic correlation than the heritability of the trait.

## Sex-linked ART locus

We assume XY sex determination for simplicity, but the results of these simulations should be equally applicable to species with ZW sex determination. In the case of X-linkage, the results are the same as for an autosomal ART locus, as long as it is possible to produce families with a mix of ARTs. For a male-limited X-linked polymorphism, heterozygote female carriers will produce mixed broods, and for a dominant female-limited X-linked polymorphism, mixed broods can be produced either by heterozygote mothers mating males carrying the recessive allele, or by homozygote mothers who mate with males carrying the opposite allele. Given that we assume that the trait and the morph locus are unlinked, results for an X-linked locus are therefore the same as for an autosomal locus since an individual’s morph assignment is independent of its breeding value for the trait. In practice, estimates will probably often be lower for X-linked loci compared to autosomal loci because of the difficulty in obtaining large numbers of offspring of each ART from the same family (Figure 1).

For a Y-linked locus, it will never be possible to produce full siblings of different ARTs, which means that some sort of maternal half-sib design is necessary for estimating inter-tactical genetic correlations. We therefore assumed a design where each mother is mated to two males, one from each ART. In this case, the family means method of estimation will consistently underestimate the genetic correlation due to the lower level of relatedness between siblings of different ARTs, so we also analysed simulated Y-linked datasets using MCMCglmm (Hadfield 2010). MCMCglmm analyses were run with a modified inverse Wishart prior, and assume that 50% of the variance in the trait is genetic in origin (Mousseau and Roff 1987). The number of iterations was 11000 with a burn-in period of 1000 and thinning interval of 50. This produced a reasonable autocorrelation in the trial runs that were carried out. The effective sample size is rather small with this combination of parameters, but was retained in the interest of saving time. Because MCMCglmm analyses are considerably more time-consuming to run than calculating the correlation based on family means (since each run involves a large number of permutations), the MCMCglmm analyses were carried out on 50 simulated datasets for each parameter combination, instead of 1000, resulting in wider confidence intervals for the MCMCglmm estimated compared to the family means estimates (Figure 2).

**Figure 2:**
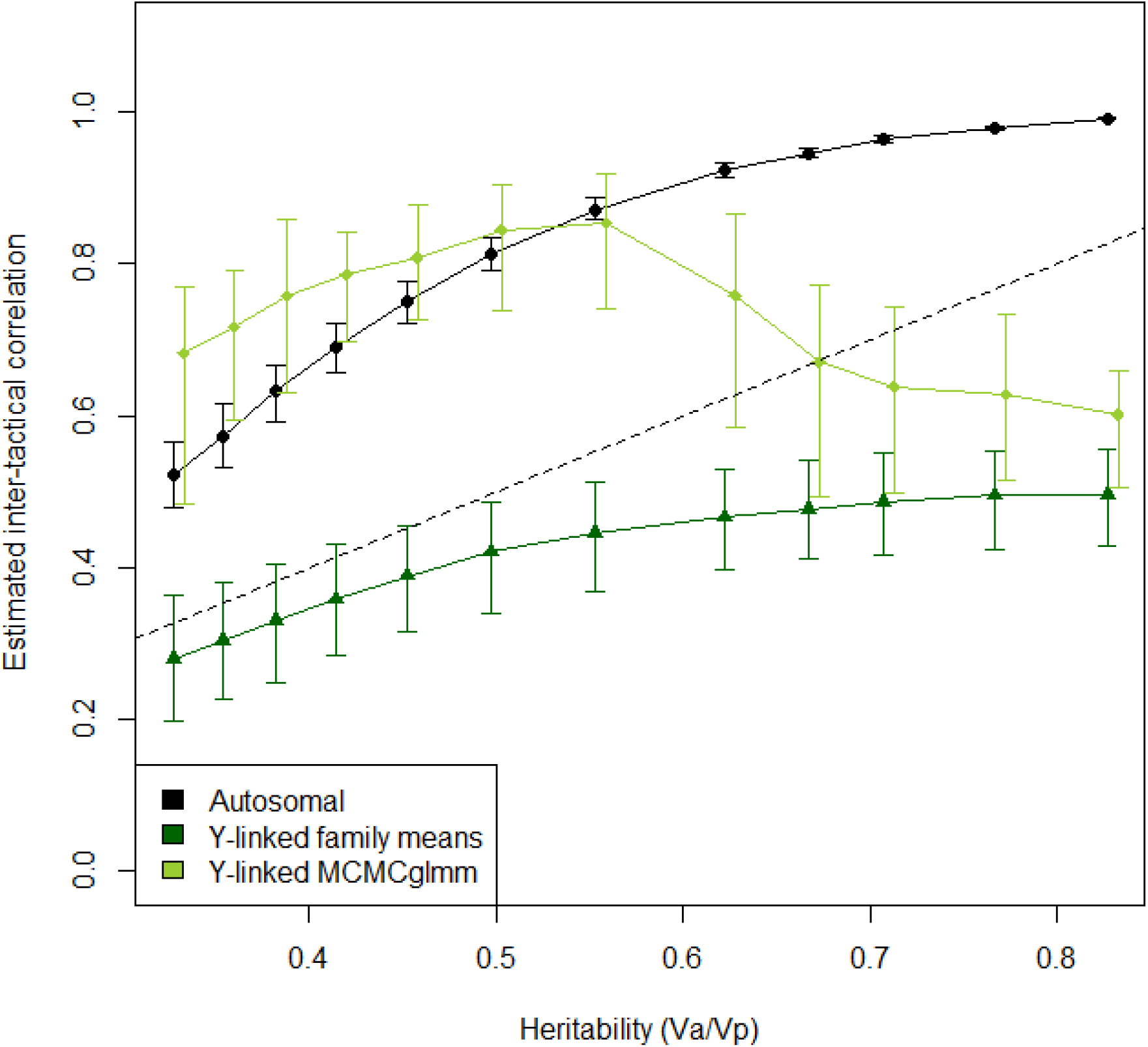
Estimated inter-tactical genetic correlation for a quantitative trait dependent on its heritability, in the case of a Y-linked ART locus. Baseline values for the autosomal case are included for the sake of comparison. Error bars denote 95% confidence intervals and points are jittered to avoid overlap. Raw family means values are approximately half that of the autosomal case, and confidence intervals are wider. MCMCglmm performs well when the prior accurately reflects the true additive genetic variance (see main text for details). Points falling above the dashed 1:1 line have a higher estimated genetic correlation than the heritability of the trait.

As expected, uncorrected inter-tactical correlations estimated from family means were about half the magnitude for Y-linked tactics compared to autosomal ones, due to the half-sib breeding design. Although accuracy of the estimates can be improved by multiplying the calculated correlation by two to take reduced relatedness into account, precision will still be considerably lower (i.e. high confidence intervals) for Y-linked tactics compared to autosomal tactics. The MCMCglmm analysis was more effective in producing estimates for Y-linked tactics that were similar to those for an autosomal locus, at least for lower heritability values (Figure 2). The decrease in estimates at high heritabilities (>0.6) is likely a side effect of the prior which assumes a heritability of 0.5. This suggests that unless information about the heritability of the quantitative trait is incorporated into the analysis, intertactical genetic correlations may be inaccurately estimated even when using MCMCglmm.

## Developmentally plastic ARTs

In the case of developmentally plastic ARTs, which tactic an individual becomes is determined by a phenotypic switching threshold. In many species, this is determined by body size, which is usually heritable, but also tends to reflect overall condition. Following Buzatto *et al.* (2015), we assumed that the switching threshold is genetically fixed in the population and the same as the population mean. We also assumed that the trait that determines the switching threshold is also the one for which we wish to estimate the intertactical genetic correlation. In practice, this means that only families with a breeding value relatively close to the population mean for the quantitative trait will produce offspring that belong to both ARTs, unless environmental manipulations make it possible to influence the developmental trajectory (we chose not to simulate this sort of manipulation for the sake of simplicity). We chose this approach because it is likely to be the least favourable scenario for detecting intertactical genetic correlations; if the trait of interest is not the trait that determines the switching threshold, then it should be much easier to obtain mixed families.

If we take body size as an example, then in this scenario, all males that are below the population mean body size at the developmental decision point become the “small” ART (i.e. sneaker or minor males) and all males that are above the population mean body size at the developmental decision point become the “large” ART (i.e. courter or major males). We found that the estimated inter-tactical genetic correlation is lower for developmentally plastic tactics compared to an autosomal ART locus, but that the estimates are still always significant as long as the heritability of the quantitative trait is moderately high (>0.3; Figure 3, dark blue triangles). The confidence interval for the estimates is also higher, which is likely a result of decreased effective sample size since not all families will produce mixed broods.

**Figure 3:**
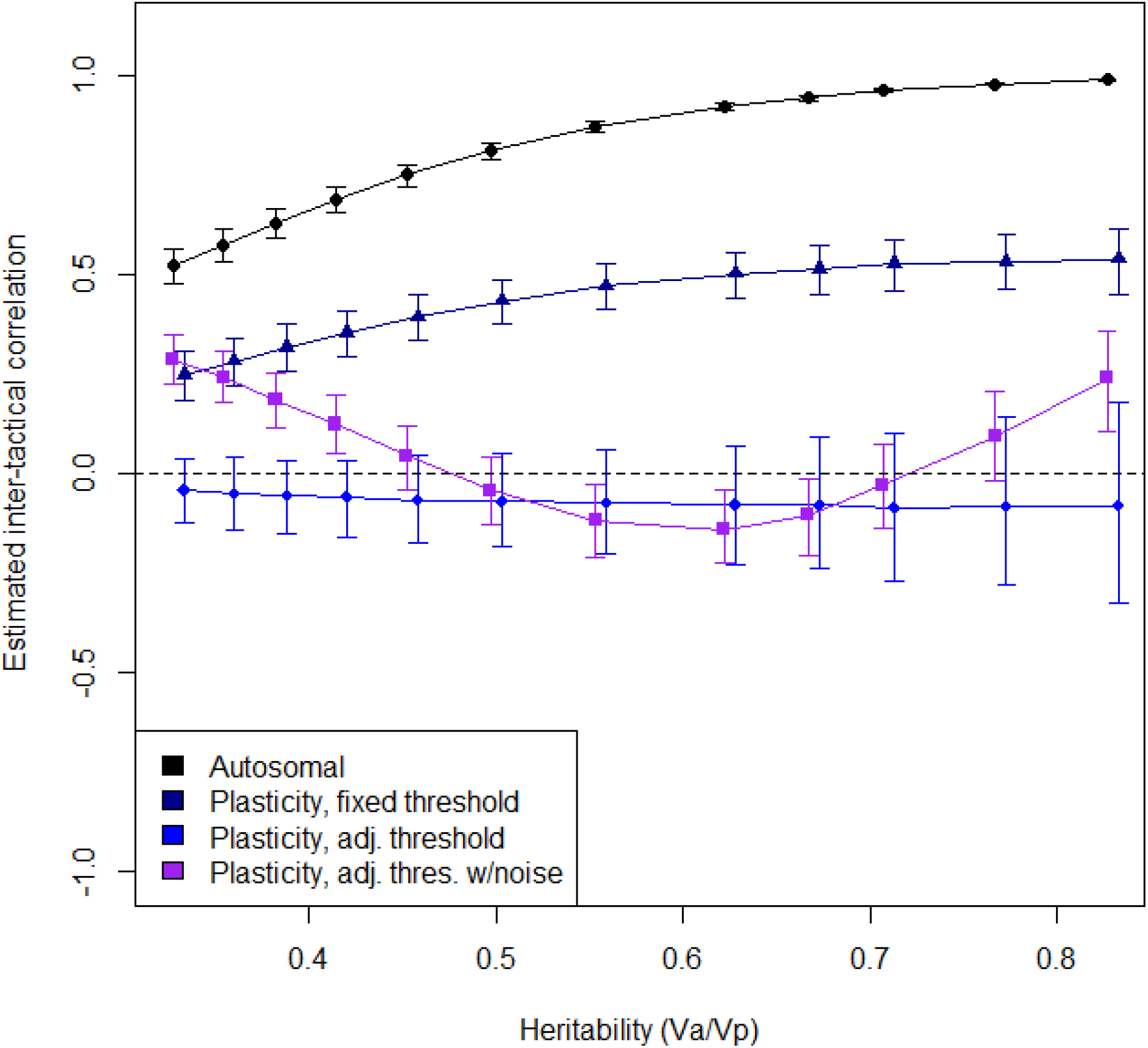
Estimated inter-tactical genetic correlation for a quantitative trait dependent on its heritability, in the case of developmentally plastic ARTs. Baseline values for the autosomal case are included for the sake of comparison. Error bars denote 95% confidence intervals and points are jittered to avoid overlap. If the switching threshold is genetically fixed in the population, then estimates are lower and more variable overall, but follow a similar pattern as in the autosomal case. If there is genetic variation for the switching threshold, then estimates often include zero (indicated by the dashed line) even though the true underlying genetic architecture is a perfect correlation.

We predict that under intra-locus tactical conflict, it would be advantageous for developmentally plastic ARTs to evolve the ability to adjust the switching threshold to match the value of the quantitative trait that determines the developmental trajectory. For example, a family with genes for becoming large will tend to produce particularly large “small” males, which may be inefficient at obtaining sneaky matings. Similarly, a family with genes for becoming small will tend to produce particularly small “large” males, which may likewise be inefficient at courting females. A family with genes for low values of the switching threshold trait should therefore adjust the threshold upwards if possible (Figure 4), and vice versa for families with high values of the trait.

There are multiple ways in which the switching threshold could be adjusted. We examine two main types of adjustment, including or excluding the effect of developmental noise. It is not our aim here to explore all possible means of adjusting the switching threshold – although there has been much speculation that species can harbour genetic variation in switching threshold, demonstrating that this is not trivial, since it requires raising similar genotypes across a range of environmental factors. This means that data on what sort of switching adjustment patterns may exist is sparse (Taborsky 2017). We therefore chose to examine two simple types of adjustment scenarios to see if and how threshold adjustment can alter the estimate of the inter-tactical genetic correlation.

If developmental noise is not taken into account when adjusting the switching threshold, then a simple rule is to move the threshold upwards or downwards the same amount as the family deviation from the population mean. For example, a family with a breeding value 0.5 standard deviations above the mean would move its switching threshold to 0.5 standard deviations below the mean, ensuring that only individuals with a trait value that is 1 population standard deviation below the family mean will develop into the “small” tactic (Figure 4). We found that this type of adjustment results in universally low and non-significant estimates of the inter-tactical genetic correlation, since the adjustment cancels out the effect of underlying genetic differences in the quantitative trait (Figure 3, light blue diamonds). Changing the magnitude of the adjustment (e.g. by multiplying the family deviation from the population mean by a fixed constant) alters the slope of the relationship between the estimate and *V_a_/V_p_*, but the general conclusions hold (Figure S1).

**Figure 4:**
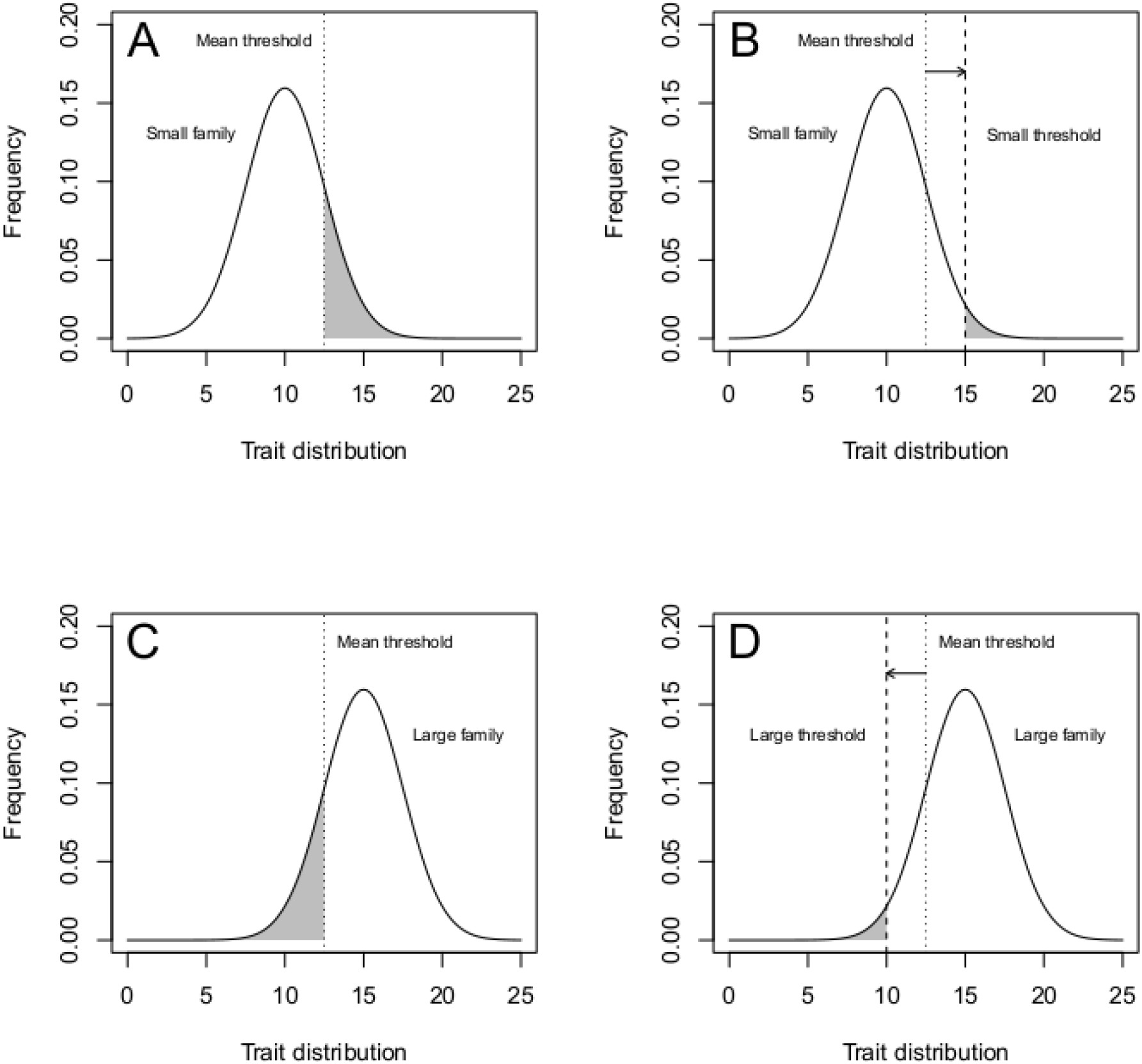
In a family of small individuals, individuals above the switching threshold will develop as “large” (i.e. major/courter, shaded area below the curve) males but be at the lower end of the size distribution for this ART (A). It would therefore be potentially advantageous for males in this family to adjust their switching threshold upwards (B). Conversely, in a family of large individuals, individuals below the switching threshold will develop as “small” (i.e. minor/sneaker, shaded area below the curve) males but be at the higher end of the size distribution for this ART (C). It would therefore be potentially advantageous for males in this family to adjust their switching threshold downwards (D). Figure partially adapted from Buzatto et al. (2015).

When developmental noise is not taken into account, as in the scenario above, this means that families with extreme values of the focal trait will not produce mixed broods unless the developmental noise parameter is very large. We therefore decided to explore what would happen if the switching threshold was adjusted relative to both the family breeding value and the degree of developmental noise. In this scenario, the threshold adjustment was scaled according to the developmental noise, such that a family with a breeding value 0.5 standard deviations above the population mean will move its threshold 0.5 within-family standard deviations (i.e. the developmental noise parameter) below the population mean. This means that the magnitude of the adjustment increases as the developmental noise increases, which we feel is a reasonable approach. The greater the uncertainty in offspring phenotype, the more scope for adjustment is needed. We found that this scenario caused estimates of the inter-tactical genetic correlation to fluctuate from negative to positive, depending on the heritability of the trait (Figure 3, purple squares). Again, changing the magnitude of the adjustment (e.g. by multiplying the threshold displacement from the family mean by a fixed constant) moves the location of the inflection point of the relationship between the estimate and *V_a_/V_p_*, but the general conclusions hold (Figure S2).

## Conclusions

Genetic correlations across ARTs is an essential criterion for intralocus tactical conflict to be a constraint on the evolution of ARTs. Our results suggest that such conflict is possible for developmentally plastic tactics, which is consistent with the detection of genetic correlations in some recent studies (Pike et al. 2017; Buzatto et al. 2018). In addition, we suggest that different methods are needed to detect genetic correlations across developmentally plastic ARTs, and that even for genetically influenced ARTs, the methods that can be applied will determine the extent to which real genetic correlations can be detected.

Inter-tactical genetic correlations are potentially more difficult to measure in developmentally plastic ARTs, unless experimental manipulations of conditions influencing the switch are possible (e.g. rearing soapberry bugs under a wide range of food and conspecific density conditions that influence a wing polyphenism, Fawcett et al. 2018; manipulated pheromones in mites to produce high status scramblers and low status fighters, Michalczyk et al. 2018). Estimates of genetic correlations were also lower for developmentally plastic tactics (at least using the family means method), meaning that non-significant results might be likely even if there is a true correlation. If there is an ability to adjust the switching threshold, all estimates become suspect. Plausible adjustment mechanisms include thresholds that are regulated by transcriptional switches. Gene-expression differences associated with polyphenic morphs have provided evidence for transcriptional switches, e.g. differences in the expression of genes encoding insulin signaling components, which alters the reaction norm for the influence of nutrition on a wing polyphenism in the soapberry bug (Fawcett et al. 2018), and additional studies reviewed by Projecto-Garcia et al. (2017). In the case of male-limited ARTs, the maternal focal trait value could also be a source of information about the family breeding value. This means that phenotypic and maternal effects cues could be combined to decide how to adjust the switching threshold (McNamara et al. 2016). Finally, in cases where the ARTs are Y-linked, half-sib breeding designs are necessary (as mixed families are not possible), and inter-tactical correlations estimated from family means will be more difficult to detect compared to the genetic correlations across autosomal tactics. In this case, experimental tests such as experimental evolution may be the only way to conclusively confirm or deny the existence of an inter-tactical genetic correlation.

In summary, substantial inter-tactical genetic correlations and therefore IATC are clearly possible for developmentally plastic tactics. However, the study of these correlations will require further understanding of both the genetic and environmental influences on the underlying thresholds.

---

**Figure S1:**
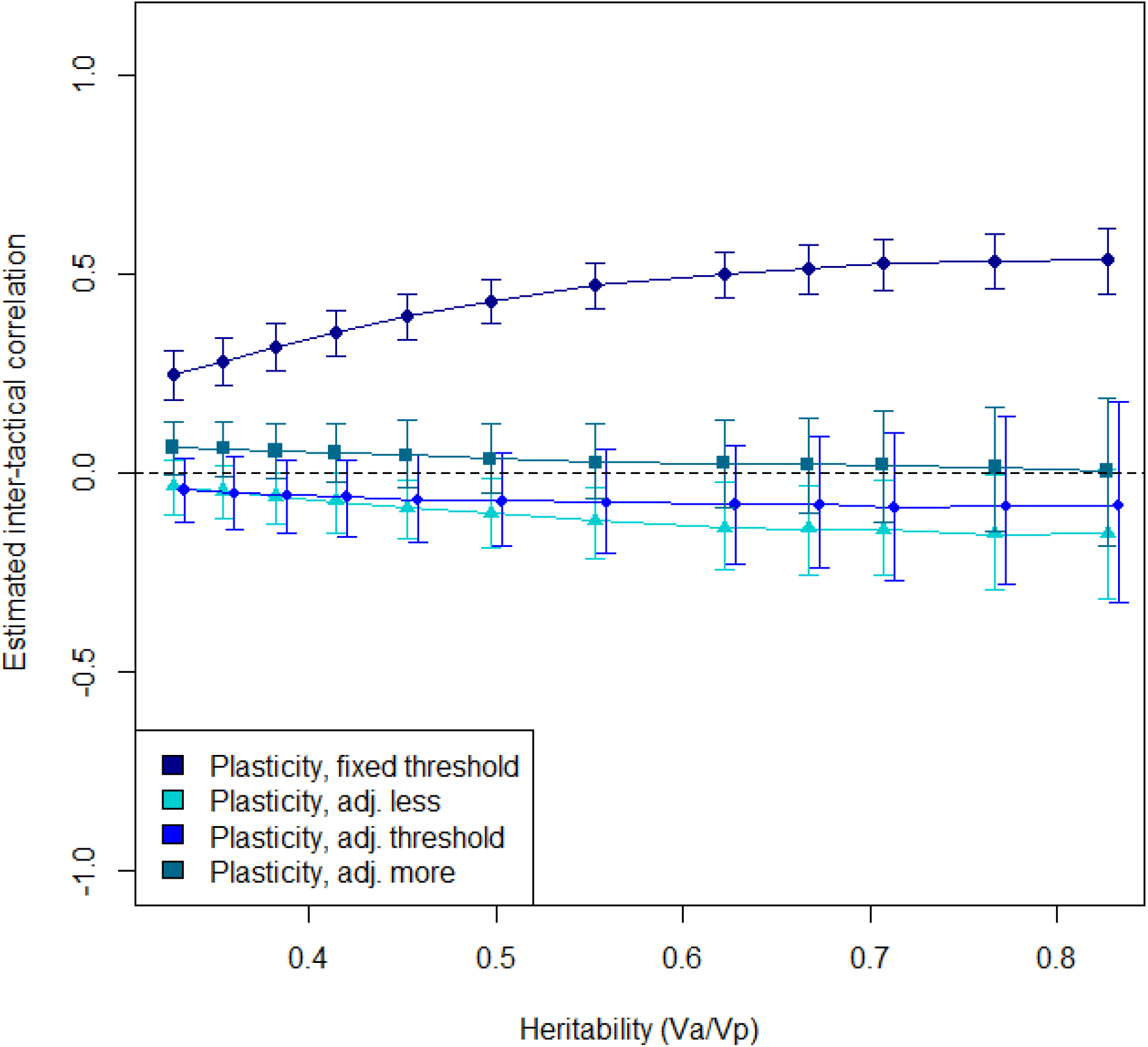
Estimated inter-tactical genetic correlation for a quantitative trait dependent on its heritability, in the case of developmentally plastic ARTs where development noise is not taken into account. Baseline values for the fixed switching threshold case are included for the sake of comparison. Error bars denote 95% confidence intervals and points are jittered to avoid overlap. Multiplying the adjustment value by a fixed constant (here 20% lower or higher, i.e. 0.8 and 1.2) changes the slope of the relationship but not the qualitative effect; estimated genetic correlations are still low and usually not significantly different from zero.

**Figure S2:**
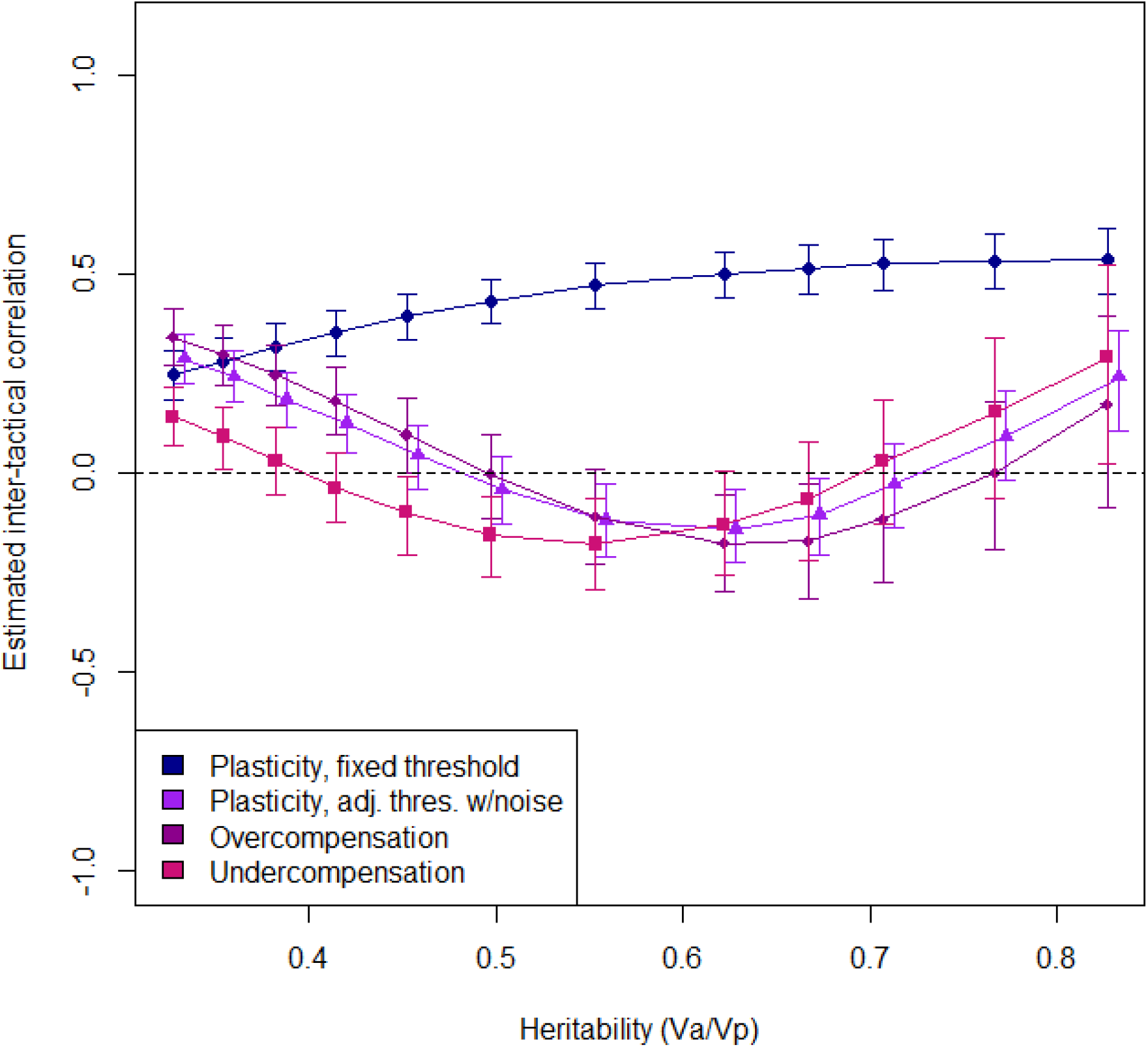
Estimated inter-tactical genetic correlation for a quantitative trait dependent on its heritability, in the case of developmentally plastic ARTs where development noise is taken into account. Baseline values for the fixed switching threshold case are included for the sake of comparison. Error bars denote 95% confidence intervals and points are jittered to avoid overlap. Multiplying the adjustment value by a fixed constant (here 20% lower or higher, i.e. 0.8 and 1.2) changes the inflection point but not the qualitative effect; estimated genetic correlations are still low and point estimates vary from negative to positive depending on the heritability of the trait.

