## Supplementary figures and images for "Genetic correlations across genetically determined and developmentally plastic alternative reproductive tactics"

### Supplemental figure 1

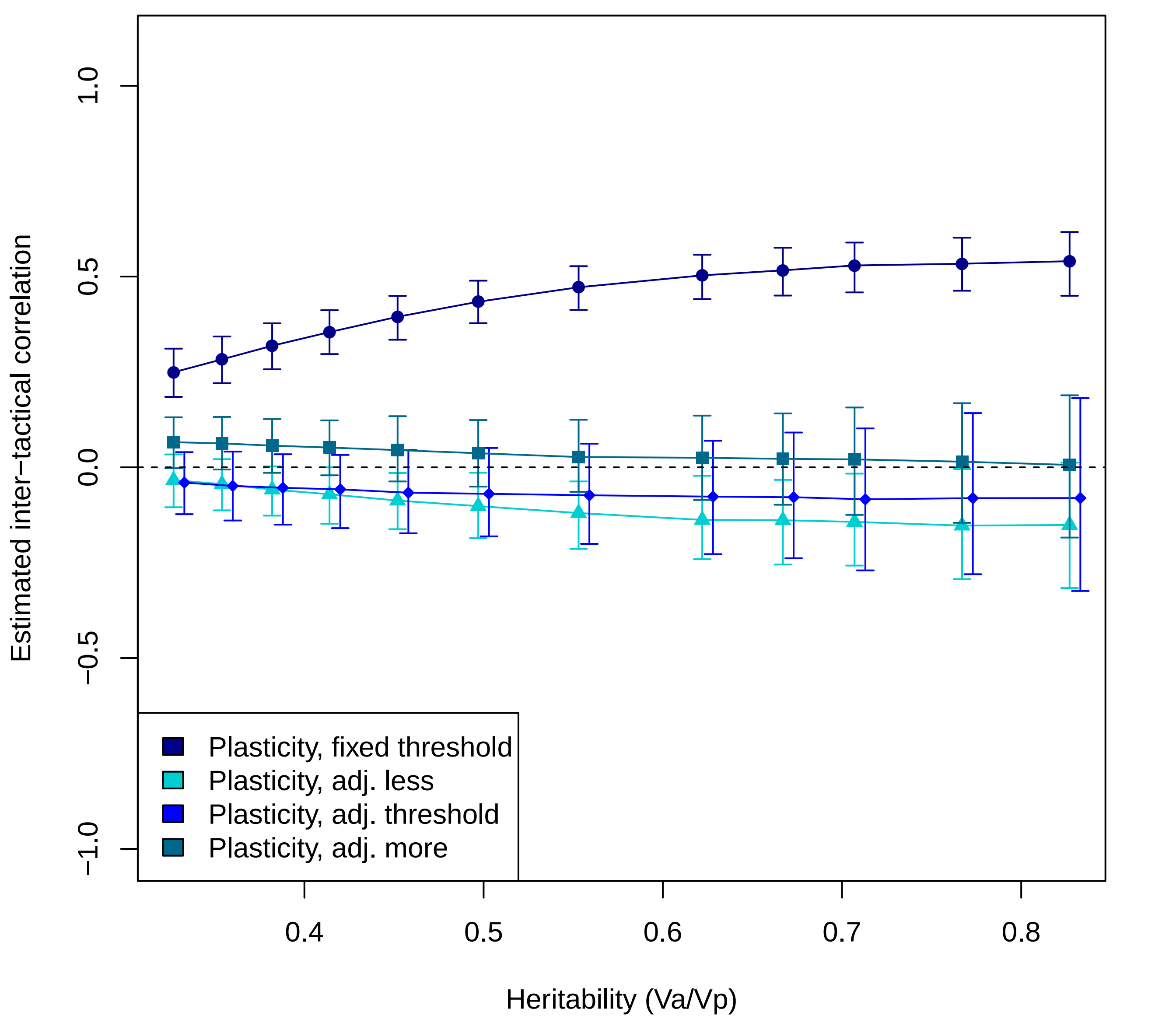

### Supplemental figure 2

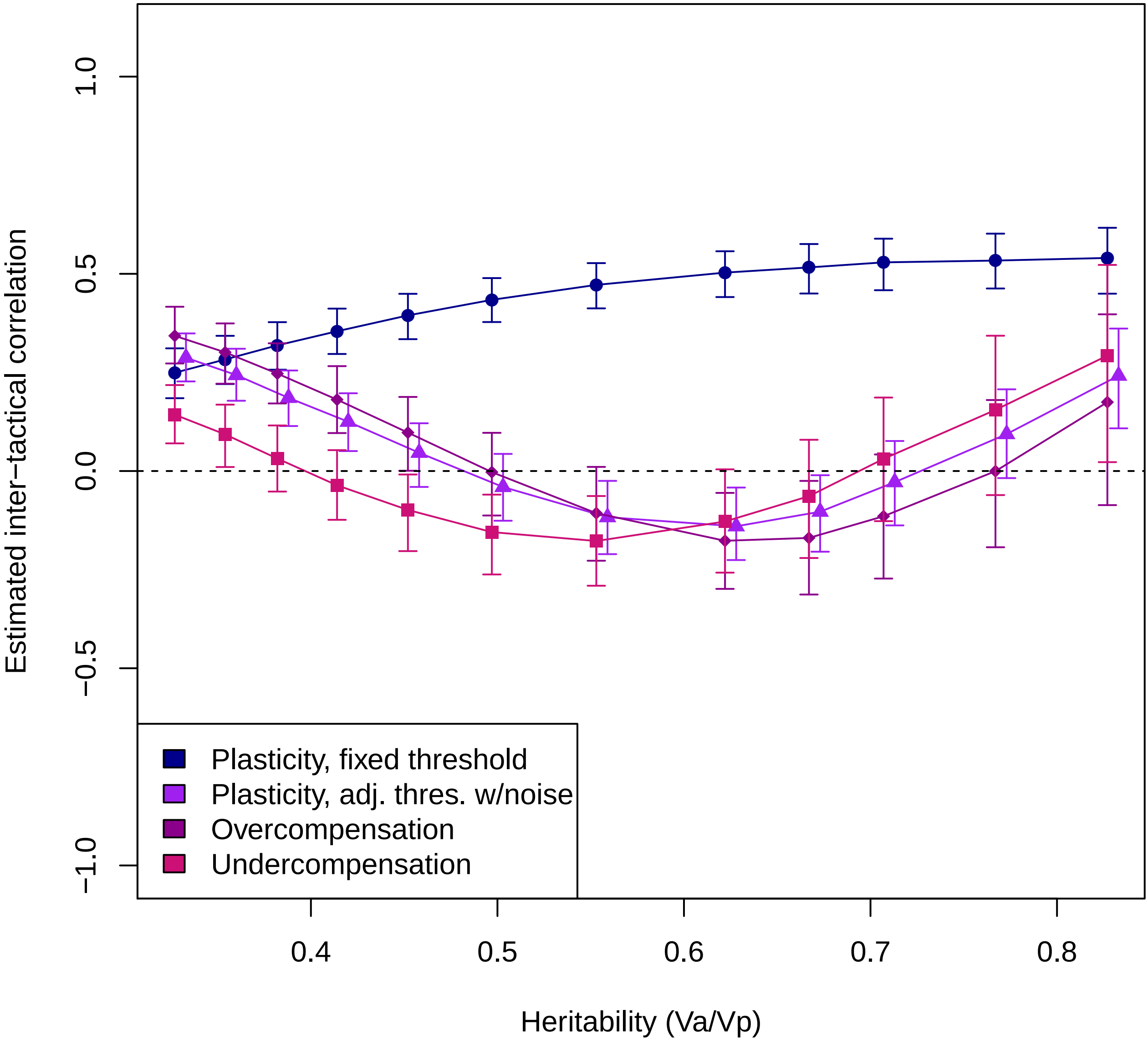
